# Aquaporin-4–Specific T Cell Responses in NMOSD Revealed by mRNA-Engineered Dendritic Cells

**DOI:** 10.64898/2026.06.05.730365

**Authors:** Verena Pauly, Maria Hastermann, Friedemann Paul, Craig C. Garner, Sara Cestari, Dietmar Schmitz, Enrico Klotzsch, Nadine Strempel

## Abstract

Neuromyelitis optica spectrum disorder (NMOSD) is an autoimmune disease of the central nervous system characterized by loss of immune tolerance to the water channel aquaporin-4 (AQP4). Current therapies do not specifically restore AQP4 tolerance or selectively suppress antigen-specific immune responses. Reprogramming patient’s own dendritic cells (DCs) with mRNA to induce tolerance represents a promising therapeutic strategy. A key prerequisite for this approach is generating mRNAs encoding disease-relevant autoantigens recognized by the patient’s immune system.

Here, we generated recombinant mRNAs encoding human AQP4 and evaluated T cell responses to autologous DCs transfected with AQP4 mRNA in eight NMOSD patients and ten healthy controls. Transfected DCs were co-cultured with autologous T cells and stimulated twice. The assay detected robust AQP4-specific T cell response in a patient with recent disease activity who was not receiving immunosuppressive therapy, demonstrating its ability to identify clinically relevant autoreactive T cell responses.

These findings establish the feasibility of an mRNA-based antigen-specific T cell assay for studying NMOSD pathogenesis, stratifying patients by T cell involvement, and supporting development of personalized tolerance-inducing therapies.

**Summary Sentence:** Recombinant mRNA encoding human aquaporin-4 (AQP4) was introduced into dendritic cells from NMOSD patients, enabling the detection of antigen-specific T cell responses, with the strongest activation observed in a patient with recent disease activity, highlighting the potential of this approach for immune monitoring and for understanding the role of T cells in the pathogenesis of NMOSD.

## Introduction

Neuromyelitis optica spectrum disorder (NMOSD) is a severe, autoimmune inflammatory and relapsing autoimmune disease of the central nervous system with oftentimes poor prognosis^**1**^. In ≥ 80% of cases, autoantibodies against the astrocyte water channel aquaporin-4 (AQP4) are detectable. A loss of tolerance to AQP4 leads to the formation of AQP4-specific autoantibodies in the periphery. These autoantibodies bind to the water channels on the astrocyte end feet, initially leading to antibody-dependent cellular cytotoxicity (ADCC) and complement-dependent cytotoxicity (CDC) followed by astrocytopathy, with subsequent loss of oligodendrocytes and demyelination of axons, particularly in the visual system and spinal cord ^**2,3**^. Consequently, affected patients suffer from motor and sensory failures as well as visual impairments, often presenting with the characteristic clinical manifestations of optic neuritis and longitudinal transverse myelitis ^**4,5**^.

At present, the pathogenesis of NMOSD is not fully understood. We and others have shown ^**6–9**^ that in addition to the B cells producing AQP4 IgG-autoantibodies, antigen-specific effector Th1 cells are also involved. This was shown e.g. through a generalized upregulation of T cell activity, as well as AQP4 specific T cells detectable after stimulation with AQP4 peptide ^**6,8,10**^. However, the processes leading to the loss of immune tolerance to AQP4 remain elusive. Central to understanding NMOSD pathogenesis is exploring how tolerance to AQP4 is disrupted and what triggers the subsequent cascade of events. A recent paper showed that AQP4 is expressed in B cells in a CD40-dependent (but autoimmune regulator (AIRE) independent) manner and suggests that thymic B cells may contribute to tolerance against a range of germinal center-associated antigens, including disease-relevant autoantigens like AQP4. These B cells are also capable of presenting their endogenous AQP4 to T cells bearing an AQP4-specific T cell receptor (TCR) ^**11**^. Intriguingly, bacterial infection and molecular mimicry of peptides between human AQP4 and bacterial AQPZ has also been found to be a possible trigger breaching tolerance ^**10,12**^.

Current treatments include the use of immunosuppressants, immunomodulators, recently FDA/EMA approved biologics based on antibodies to kill-off antibody producing B cells, as well as complement inhibitors and IL-6 blockers to blunt the immune response. However, these are not specific and have numerous and serious side effects of long-term immune suppression ^3,13^ motivating the search for therapies that directly and selectively restore tolerance to AQP4.

One potential strategy to re-establish specific immune suppression is through the *ex vivo* reprogramming of peripherally circulating dendritic cells (DCs) ^14^. DCs are key regulatory players in the immune system, orchestrating a delicate balance between immune activation and tolerance maintenance. Through their ability to capture, process, and present antigens in either a pro-inflammatory or homeostatic environment, DCs can either stimulate the immune response by activating effector T cells and antibody producing B cells or promote tolerance by inducing regulatory T cells specific to the presented peptides ^15,16^. Importantly,the immunomodulatory capacity of DCs represents a significant opportunity to manipulate the immune system in an antigen-specific (Ag-specific) manner. This is best exemplified by previous studies demonstrating the feasibility of generating an immune response to cancer antigens by reprogramming DCs ^17–19^.

*Ex vivo*-reprogrammed tolerogenic DCs hold potential for restoring Ag-specific tolerance in autoimmune diseases. Efficient delivery of antigens to DCs for peptide processing remains a challenge. Approaches such as targeting DC-specific receptors ^20,21^, plasmid nucleofection ^22^ and employment of viral vectors ^23^ have demonstrated efficiency but are constrained by a limited peptide repertoire. Peptides are commonly used for Ag-specific stimulation, particularly in animal models ^6,9,11^. However, the pathogenicity of AQP4 *in vivo* remains low due to tolerance mechanisms. Even when some of these mechanisms were eliminated ^11^, proper disease development was not observed, likely due to suboptimal peptide characteristics such as length, tagging, and modifications, which differ from the processing of the complete protein by cells. Furthermore, even with MHCII matching for the respective peptides, a strong phenotype in animals was not achieved ^6^. Strategically, this issue might be addressed by expressing recombinant full-length AQP4 mRNA that includes an endo-lysosomal targeting element and is HLA-independent; recognizing that individual protein processing varies among people. In addition, using this approach could generate *ex vivo* autologous tolerogenic DCs before returning them to patients.

In the current study, we have explored the feasibility of generating an endo-lysosomal targeted version of human AQP4 (AQP4*) that when expressed as an mRNA in DCs is processed in a manner that is recognized by T cells from patients with active NMOSD. Our results show that our recombinant AQP4* is not only sorted through the endo-lysosomal system in mDCs but also when expressed in DCs, can elicit an immune response in T cells from healthy individuals upon co-expression with an immune activating molecule. Importantly, AQP4*-mRNA expression alone was found to robustly activate Ag-specific (CD4^+^ CD154^+^) T cells in a NMOSD patient with a positive AQP4-antibody titer and relapses in the past 31 months, but not in those patients receiving immune-suppressive drugs or disease activity more than 6 years ago. These results suggest that our AQP4*, expressed *via* its mRNA, can be properly processed and thus potentially used to explore in detail the role of T cells in NMOSD pathogenesis, as a diagnostic tool to identify patients with T cell involvement as well as the development of mRNA-based Ag-specific therapies for NMOSD.

## Materials and Methods

### Cell Media

CD14^+^ cells (monocyte derived dendritic cells (moDCs)) were cultured in X-Vivo 15 Media (Lonza, 02-060Q) with 800U GM-CSF and 250U IL-4 (PeproTech, AF-300-03, AF-200-04).For T cell feeding R10 Media was used adapted by Cimen Bozkus et al (2021) ^24^: RPMI 1640 Media was supplemented with HEPES (10mM), Gentamycin (0,1mg/mL), GlutaMax (1x) and Human Serum (10%) and supplemented with IL-2 (Gibco, PHC0027), IL-7(R &D Systems, BT-007-GMP-025) and IL-15 (PeproTech, 200-15).

T cells (CD14 Flow through) were cultured in RPMI Media supplemented with 1% Penicillin/ Streptomycin and 10% FCS for one week before co-culture with autologous mDCs.

### Patient and Healthy Donor Samples

Experiments were performed with blood from 10 healthy individuals provided by Zentrum für Transfusionmedizin und Zelltherapie Berlin (ZTB), Charité Campue Mitte using leukocyte reduction filters. Venous blood samples were also collected from eight AQP4-IgG-positive diagnosed patients from the BERLimmun cohort ^25^, fulfilling the current diagnostic criteria for NMOSD (See Table 1) ^26,27^.

**Table 1:** Clinical Data/information of NMOSD patients. Therapy = only immunomodulatory therapy, RTX = Rituximab, all data are collected at the time of blood collection, *=before blood collection,

| Pat No | Sex | Age | Therapy | AQP4-IgG titer | Total relapses | Last relapse | EDSS Score |
| --- | --- | --- | --- | --- | --- | --- | --- |
| 1 | F | 57 | None | 1:320 | 3 | 2021 | 3.0 |
| 2 | F | 49 | None since 2017 | No value (last positive 2014) | 1 | 2014 | 2.0 |
| 3 | F | 27 | RTX (since 15.09.23) | 1:32 | 4 | 2016 | 3.5 |
| 4 | F | 45 | RTX | 1:32 | 4 | 2015 | 2.0 |
| 5 | F | 62 | RTX | neg | 3 | 2013 | 5.0 |
| 6 | F | 68 | Inebilizumab | 1:32 | 2 | 2017 | 4.0 |
| 7 | F | 60 | RTX | ? | 7 | 2010 | 7 |
| 8 | F | 59 | none | 1:320 | 2 | 2011 | 4.5 |

### Ethics Statement

This study was approved by the Ethics Committee of the Medical University of Berlin (license EA4/099/20 and EA1/041/14 /T regs).

### Design of Expression Vectors Encoding AQP4*

Human AQP4 (AQP4) is processed into two isoforms ^28^. The one lacking its amino-terminal 23 amino acid residues, AQP4-M23, forms Orthogonal Arrays of Particles (OAPs), which facilitates a better auto-antibody binding compared to M1 isoform ^28,29^. We therefore designed our recombinant AQP4 according to the more OAP prone isoform to start at methionine 23 (Fig. 1A-C). We furthermore added a Flag epitope in loop C allowing visualization by immunofluorescence microscopy resulting in AQP4-M23-flag-wt (AQP4-wt). The flag epitope was inserted in the middle of the construct between Valine141 and Valine142, following the approach of Verkman et al. (2012) and Crane et al. (2008) ^30,31^ (Fig. 1C). These former studies confirmed that this specific location does not interfere with the formation of orthogonal arrays of particles (OAPs). We chose this position because it ensures that AQP4 would be endo-lysosomally degraded, which is crucial for our experimental design (see below). To facilitate peptide processing and presentation of AQP4 peptides in the context of MHCII, we also tagged a version of AQP4-M23-flag with the endo-lysosomal targeting signaling sequence from lysosome-associated membrane protein-3 (LAMP3), a dendritic cell specific endo-lysosomal protein ^30,32^. This targeting sequence is found at the C-terminal cytoplasmic tail of LAMP3 and is functional only if situated proximal to endosomal membranes ^33^. Unfortunately, AQP4 cannot be appropriately tagged due to its long cytoplasmic C-terminus. To overcome this challenge, we appended the C-terminus of AQP4 with two additional transmembrane domains (TMD). One of these was from the synaptic vesicle protein VAMP2 and the second from the last TMD from LAMP3 containing the endo-lysosomal targeting element, 11 amino acids from its TMD-cytoplasmic interface. This arrangement, creating AQP4-M23-flag-Lamp3 (AQP4*) (Fig. 1C), is predicted to place the LAMP3 endo-lysosomal targeting element on the cytoplasmic face of endosomes, where it is then recognized by the lysosomal sorting system ^33^.

**Figure 1.**
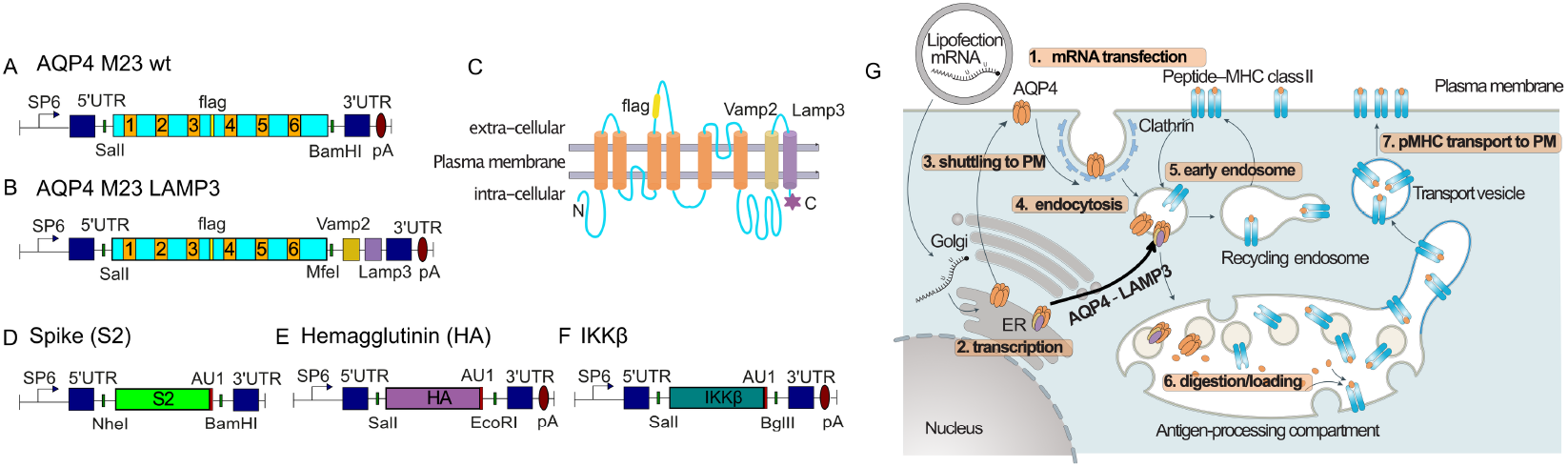
Engineering of AQP4 mRNA constructs and strategy for enhanced MHC class II antigen presentation. (A) Schematic representation of the wild-type AQP4 M23 mRNA construct containing SP6 promoter, 5′UTR, coding sequence, introduced FLAG tag, and 3′UTR followed by a poly(A) tail. Relevant restriction sites are indicated. (B) Modified AQP4 M23 construct fused to VAMP2 and LAMP3 targeting sequences to promote trafficking to endo-lysosomal compartments (AQP4–LAMP3). (C) Predicted membrane topology of the AQP4–LAMP3 fusion protein, showing the six transmembrane domains of AQP4, extracellular FLAG tag, and C-terminal fusion to VAMP2/LAMP3 domains directing intracellular sorting. (D–F) Control mRNA constructs encoding SARS-CoV-2 Spike S2 subunit (D), Influenza hemagglutinin (HA) (E), and IKKβ (F), each flanked by untranslated regions (UTR) and poly(A) tail (pA) as indicated, AU1 serves as tag for Immunofluorescence detection. (G) Model of intracellular processing following lipofection-mediated mRNA delivery. Transfected mRNA is translated and AQP4 or AQP4–LAMP3 proteins traffic to the plasma membrane, followed by clathrin-mediated endocytosis and routing through early and recycling endosomes. The LAMP3 fusion promotes targeting to antigen-processing compartments, where proteins undergo proteolytic degradation and peptide loading onto MHC class II molecules. Peptide–MHC complexes are subsequently transported to the plasma membrane for antigen presentation to CD4^+^ T cells.

### Design of Additional mRNA Expression Vectors

As control antigens, we designed and utilized two engineered mRNAs encoding either the influenza-hemagglutinin protein or the SARS-CoV-2 SPIKE protein. The HA-mRNA (HA) encodes the glycoprotein that envelops the influenza virus, based on the H1N1 variant (A/Guangdong-Maonan/SWL1536/2019) (H1N1) (Fig. 1E). The S2-mRNA (S2) encodes the S2 region of the SARS-CoV-2 spike protein (Fig. 1D), which shows a preference for eliciting T cell responses ^34^. The S2 domain, representing the stalk region of the SPIKE protein, is highly conserved ^35^ and recognized by both MHCII and MHCI ^36^. As an immune activator,we created a mRNA encoding a constitutively active (CA) stabilized mutant of IKKβ (Fig. 1F), as reported by Pfeiffer et al ^37^. Specifically, to activate IKKβ, we exchanged serine residues within its activation centers with phospho-mimicking glutamic acid residues. To increase stability, we destabilized the PEST (proline (P), glutamic acid (E), serine (S) and threonine (T) enriched) region by replacing specific serine (Ser) residues in this region with alanine (Ser670, 672, 675, 679, 682, 689, 692, 695, 697, 705). Of note, IKKβ is known to play a crucial role in the activation of the NF-κB-pathway, a pivotal mediator of pro-inflammatory responses and regulator of T cell survival, activation and differentiation of inflammatory T cells as well as innate immune-cells ^37,38^.

The mRNAs for each of these proteins were designed *in silico*. To facilitate mRNA translation and stability, the coding region from each was placed between the 5’ and 3’ untranslated regions (UTRs) of hemoglobin and a Kozak translational initiation sequence was placed 5’ of the initiating methionine. This design also includes a SP6 promoter, allowing *in vitro* transcription as well as the production and purification of mRNAs encoding each of the constructs (see below). To allow the detection by immunofluorescence microscopy, each C-terminal (HA, S2, IKKβ) was tagged with an AU1 epitope (DTYRYI; BioLegend, 901901).

### Vector Construction

Plasmids pEX-A258-AQP4-M23-Flag, pEX-A258-AQP4-M23-Flag-Vamp2-Lamp3, pEX-A258-S2spike-AU1, pEX-A258-HA-AU1 and pEX-A258-CA pest IKKβ-AU1 were provided by Eurofins Genomics (Ebersberg, Germany), transformed into NEB 5-alpha Competent *E. coli* (New England *Biolabs)* and purified. Five constructs were created:

1. **AQP4-M23-Flag (AQP4-wt)**: pEX-A258-AQP4-M23-Flag was cut with the enzymes SalI and BamHI, and used to replace the mRFP coding region in the destination vector pCMV-mRFP-polyA that was also opened with SalI and BglII.
2. **AQP4-M23-Flag-Vamp2-Lamp3 (AQP4*)**: pEX-A258-AQP4-M23-Flag was cut with the enzymes SalI and MfeI, and inserted into the vector pCMV-Lamp3-polyA TSHRr was opened with SalI and EcoRI.
3. **S2:** pEX-A258-S2spike-AU1 was cut with the enzymes NheI and BamHI, and used to replace the EGFP coding sequence in the vector pEGFP-C1 was cut with NheI and BamHI.
4. **HA:** pEX-A258-HA-AU1 was cut with the enzymes SalI and EcoRI, and used to replace ethe RFP coding region in the vector pCMV-mRFP-polyA was cut with SalI and EcoRI.
5. **IKKβ:** pEX-A258-CA pest IKK-β-AU1 was cut with the enzymes SalI and BglII and used to replace ethe RFP coding region in the pCMV-mRFP-polyA vector was cut with SalI and BglII.

The resulting fragments were ligated into their respective vectors (pCMV mRFP-polyA, pCMV-Lamp3-polyA, pEGFP-C1) using T4 DNA ligase (New England Biolabs, Frankfurt Germany) after removing the mRFP or GFP coding region in the destination vector. The new vectors include a CMV promoter for expression in HEK293 cells, SP6 promoter for *in vitro* mRNA transcription (IVT) to produce functional mRNAs and except for the S2 construct, a long polyA-tail, to increase translational efficiency.

### mRNA Transcription and Testing

*In vitro* mRNA transcription (IVT) was performed with mMESSAGE mMACHINE™ SP6 Transcription kit according to the manufacturer’s protocol (Thermo Fisher, AM1340, Germany). As the S2 construct lacks an intrinsic poly-A stretch, a polyA tail was added after IVT using the poly(A) tailing kit, (Thermo Fisher, AM1350). Purity was initially assessed using agarose gel electrophoresis, before being transfected into HEK293 cells or human monocyte derived DCs (moDCs) using Lipofectamine™ MessengerMAX™ Transfection Reagent (Thermo Fisher, LMRNA001) performed according to the manufacturer’s instructions. The transfected and fixed cells were immunostained with rabbit anti-FLAG (Sigma, F7425) and mouse anti-LAMP1 (Abcam, ab25245) or mouse anti-AU1 for S2-AU1, HA-AU1, IKKβ-AU1 (biolegend, 901901) primary antibodies followed by goat anti-mouse IgG Alexa Fluor™ 488 (Thermo Fisher, A-10680) or goat anti-rabbit IgG Alexa Fluor™ 647 (Thermo Fisher, A-11008) as secondary antibodies. Hoechst 33342 was added as a nuclear stain. Proper expression of S2, HA and IKKβ in HEK293 as well as in mDCs was confirmed as shown in **Figure S1**. Images were taken with a Spinning Disc Confocal Microscope (Carl Zeiss Axio Observer.Z1 with Andor spinning disk and cobolt, omricon, i-beam laser) (Carl Zeiss, Andor) using a 40x (1.3 NA) Plan-Apochromat oil objective and an iXon ultra (Andor) camera controlled by iQ software (Andor) or a Nikon Spinning Disk Confocal under 40x (1.3 NA) objective. All images were taken with a Z stack of 5 μm with a step size of 0.5 μm.

### Overlap Analysis of Fluorescent Signals

The overlap between AQP4 variants and LAMP1 was analyzed using scatterplots generated using Python version 3.12.0 (PyCharm 2022.2.2). The main packages used for data processing were numpy version 2.1.3, scikit-image version 0.24.0 and scipy 1.14.1 for segmentation and pandas version 2.2.3. Matplotlib version 3.8.3 was used for plotting. For each image, cell regions of interest (ROIs) were defined using thresholding based on the AQP4 channel. A watershed algorithm was than applied to segment cells per each field of view. Within the thresholded regions, pixel intensities of AQP4 variant and LAMP1 intensities were plotted as a scatter plot for the center planes of the respective cells. Per segmented cell, the background plus standard deviation was calculated to define the intensity threshold per channel (using ImageJ). Number of pixels to total pixels allowed quantifying overlap as percentage and per condition.

### Spot Detection and Quantification

Spot detection analysis of AQP4-positive regions at the basal plane of segmented cells was conducted using Fiji/ImageJ using the Find Maxima function. Detected spots were marked automatically based on intensity thresholds, and results were verified manually to exclude background artifacts. The total number of detected spots was calculated for each cell, and comparisons were made between AQP4* and AQP4-wt conditions. Statistical differences were assessed using a two-tailed t-test, with a significance threshold of p < 0.05.

### Violin Plot Generation

Violin Plots comparing the number of AQP4-positive spots at the basal membrane for AQP4* and AQP4-wt were generated using Python Pycharm package. The plots visualize the distribution, median, and range of spot counts across cells and highlight statistically significant differences.

### PBMC Isolation

The strategy used in the reprogramming and testing of mDCs is shown in **Figure S3**. The isolation of human peripheral blood mononuclear cells (PBMCs) was performed with CPT collection tubes/ BD Vacutainer® CPT™ System according to the manufacturer’s protocol. PBMCs from healthy controls were isolated from leukocyte reduction filters by back flushing with PBS/EDTA/HSA followed by Ficoll gradient centrifugation. PBMCs from patients and healthy individuals were cryopreserved and thawed for restimulation of the T cells.

### Generation of Monocyte-Derived Dendritic Cells (moDCs)

CD14^+^ MicroBeads (Miltenyi Biotec, 130-050-201, Bergisch Gladbach, Germany) were used for monocyte separation from PBMCs. CD14^+^ cells were cultivated in X-Vivo 15 Media (Lonza) supplemented with 800U/ml GM-CSF and 250U/ml IL-4 (PeproTech, AF-300-03, AF-200-04) for 6 days resulting in differentiation into moDCs. Maturation of moDCs was induced for additional 24 h with TNFα, IL-1β (PeproTech AF-300-01A, AF-200-01B), IL-6 (Miltenyi Biotech, 130-093-929) and prostaglandin E2 (PGE2) (Enzo, BML-PG007-001) resulting in mDCs ^39^. The CD14 negative fraction, containing T cells, was cultivated in RPMI1648 media.

### mRNA Transfection of mDCs and Pan T cell Isolation

The mDCs were transfected at day 7 with the engineered mRNAs (AQP4-wt, AQP4*, IKKβ, HA, or S2) using Lipofectamine™ MessengerMAX™ Transfection Reagent. Autologous T cells from the cultivated flow through were harvested and MACS-sorted *via* Pan T cell isolation Kit (Miltenyi Biotec, 130-096-535) according to the manufacturer’s instructions.

### Co-culture of mDCs and T cells

Autologous T cells were added after 1.5 h to the mRNA transfected mDCs and stimulated for 20-24 h. Half of the media was exchanged every 2-3 days with R10 media.

At day 14 the T cell restimulation was repeated according to the protocol described (**Fig. S3**) to enhance the activation of rare Ag-specific T cell populations ^40^. Restimulation was performed for 20-24 h in the presence of CD40-antibody (1µg/ml, Miltenyi Biotec, 130-094-133) to capture transient CD154 expression for surface staining ^40^. Brefeldin A (2µg/ml) was added to the cells for the last 1.5 h of stimulation to block cytokine secretion.

### Immunocytochemistry Staining

To analyze T cell responses (**Fig. S3**), the cells were stained extracellularly and intracellularly, followed by flow cytometric analysis. T cells were carefully harvested and washed once with PBS. Surface staining was performed by incubating T cells with fluorochrome-conjugated antibodies for 20 minutes (**Supplementary Table 1)**. After fixation and permeabilization using Inside Fix (Miltenyi Biotec, 130-090-477) for 20 min, intracellular staining was performed using antibodies listed in **Supplementary Table 2** for 20 minutes. All staining steps were carried out in the dark at room temperature. Stained T cells were analyzed by flow cytometry using a Symphony A5 (BD Biosciences) device. Gating was done as described in **Figure S4**.

Data analysis was performed using FlowJo software (versionX.10.10.0, BD Biosciences), with appropriate compensation applied to account for spectral overlap.

### Statistics

All statistical analyses were performed using GraphPad Prism (version 10 or later). Data are presented as individual data points with median ± interquartile range (IQR), unless otherwise indicated. For comparisons between paired conditions (e.g., stimulated vs. unstimulated samples from the same donor), two-tailed paired non-parametric tests (Wilcoxon matched-pairs signed-rank test) were applied. Statistical significance was defined as *p* < 0.05. Exact *p* values are reported where appropriate. Furthermore, this was an exploratory study with no adjustment for multiple comparisons.

## Results

### Development of Endo-Lysosomal Processed Human AQP4 for Enhanced T cell Activation

Advancements in mRNA technology provide a promising platform for optimizing antigen presentation^17,18,41^. Here, mRNA-transfected DCs were shown to mimic the natural processing pathway from mRNA to peptides, presenting a broader repertoire of peptides bound to major histocompatibility complex molecules (MHC-I or MHC-II). To leverage this, we designed mRNA expression vectors encoding human AQP4 (AQP4), with features to enhance immunogenicity and facilitate detection and endo-lysosomal targeting (Fig. 1A-C). First, to enhance immunogenicity we employed a variant of AQP4 with a higher antibody binding propensity lacking its initial 23 amino-acid residues and containing an initiating translation site at methionine 23, AQP4-M23^42^. Secondly, a Flag epitope was added in the extracellular loop C for easy detection as well as a C terminal endo-lysosomal targeting element from LAMP3 (AQP4*), a DC-specific endo-lysosomal protein ^32^, previously used in cancer immunotherapy to improve antigen processing and immune responses.

To prove if mRNA can be used as a general alternative to peptides inducing antigen-specific T cell responses in humans, we also designed mRNA constructs encoding Spike (S2) and hemagglutinin (HA) (Fig. 1D, E). In addition, to overcome natural tolerance to the autoantigen AQP4, we employed a constitutively active IKKβ variant to stimulate the NF-κB signaling pathway (Fig. 1F).

The strategy for enhanced antigen presentation via the MHC class II pathway is depicted in Fig. 1G. In this approach, mRNA is introduced into matured dendritic cells (mDCs) via lipofection. The inclusion of a LAMP3 targeting signal is intended to accelerate antigen processing by directing the AQP4–LAMP3 fusion protein to the endolysosomal compartment. There, it can be efficiently loaded onto MHC class II molecules, followed by trafficking to the plasma membrane for presentation to antigen-specific T cells.

### Expression in HEK293 and Mature Dendritic Cells (mDCs)

In an initial series of experiments, we evaluated both the expression and subcellular localization of the recombinant AQP4-M23-wt (AQP4-wt) and AQP4-M23-LAMP3 (AQP4*) constructs in transfected HEK293 cells (Fig. S2). Cells were transfected either with plasmid vectors, as described in the Methods, or with *in vitro* transcribed mRNA generated using SP6 DNA-dependent RNA polymerase. Six hours post-transfection, cells were fixed and stained with FLAG antibodies to identify transfected cells and assess the distribution of the AQP4 variants. For experiments in monocyte-derived dendritic cells (moDCs), cells were transfected with the corresponding mRNAs ^43,44^.

To assess membrane localization of AQP4, we quantified AQP4-positive puncta at the basal membrane in cells expressing AQP4 constructs. Imaging revealed sparse AQP4^+^ spots in the basal plane (Fig. 2A), whereas expression of wild-type AQP4 resulted in a marked increase in puncta density consistent with its known ability to form higher-order membrane assemblies known as orthogonal arrays of particles (OAPs). (Fig. 2B). Quantification confirmed a significant elevation in AQP4 spot number at the basal membrane in AQP4-wt cells compared to control (Fig. 2C). Consistently, the total cell adhesion surface area was increased in AQP4-wt-expressing cells (Fig. 2D), indicating enhanced membrane organization and/or spreading.

**Figure 2.**
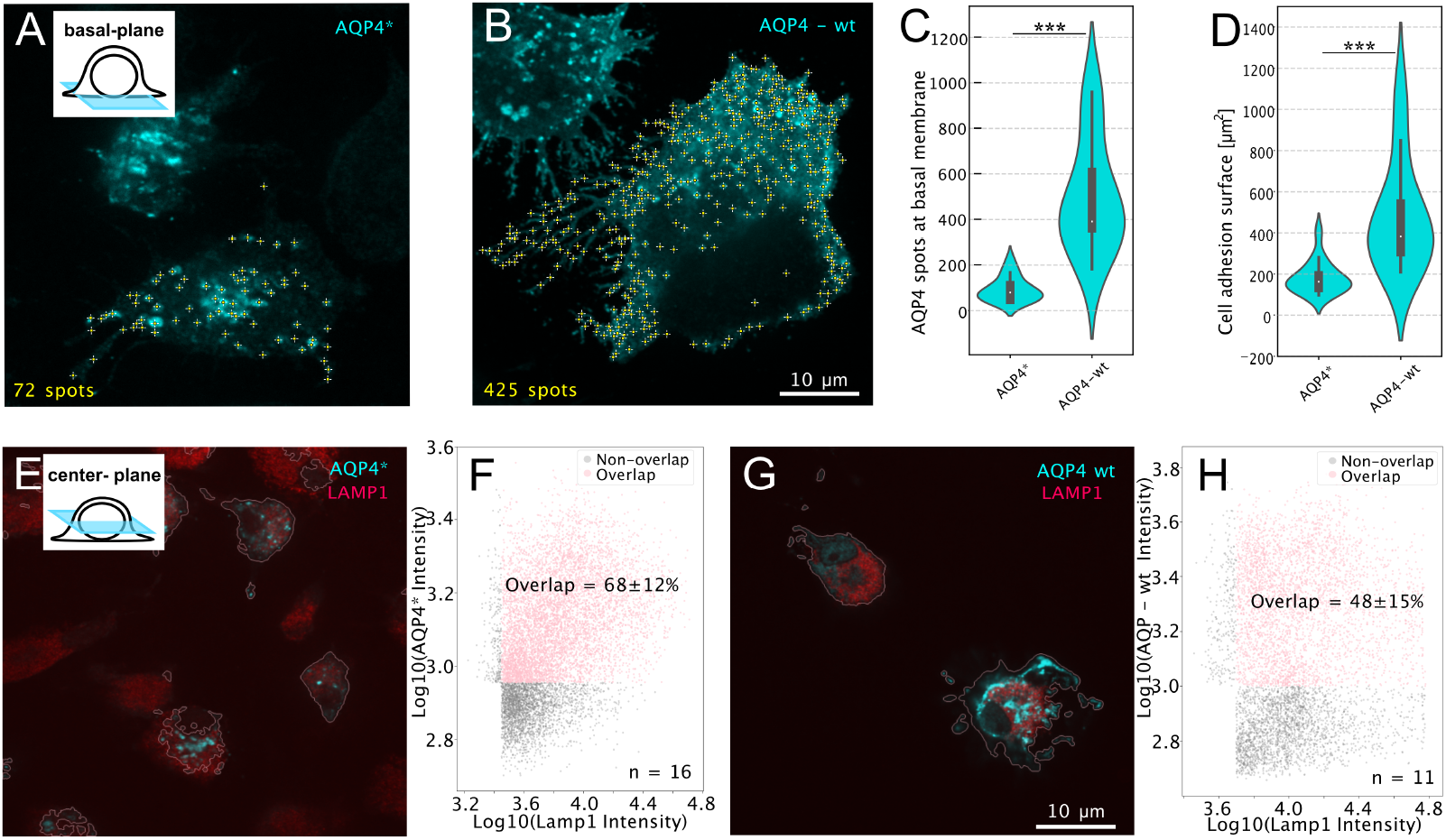
AQP4-LAMP3 targeting decreases basal membrane accumulation and enhance lysosomal co-localization. (A, B) Representative basal-plane fluorescence images of cells expressing AQP4* (A) or wild-type AQP4 (AQP4–wt) (B). Detected AQP4-positive spots are indicated (yellow crosses). AQP4–wt exhibits a marked increase in basal membrane-associated puncta compared to AQP4*. Insets illustrate imaging planes. Scale bar, 10 μm. (C) Quantification of AQP4-positive spots at the basal membrane reveals a significant increase in AQP4–wt compared to AQP4* (***P < 0.001). (D) Cell adhesion surface area is significantly increased in AQP4–wt-expressing cells relative to AQP4* (***P < 0.001). (E, G) Representative center-plane images showing co-staining of AQP4 constructs (cyan) with the lysosomal marker LAMP1 (red) for AQP4* (E) and AQP4–wt (G). Cell boundaries are outlined. Scale bar, 10 μm. (F, H) Quantitative analysis of AQP4 and LAMP1 signal overlap. Scatter plots of log10-transformed intensities indicate a higher degree of co-localization for AQP4* (68 ± 12%, n = 16 cells) compared to AQP4–wt (48 ± 15%, n = 11 cells). Overlapping and non-overlapping populations are indicated.

We next examined intracellular trafficking of AQP4 and its targeting to the endolysosomal compartment. Confocal imaging of the center plane showed substantial co-localization of AQP4 with LAMP1 (Fig. 2E). Quantitative analysis revealed a high degree of overlap between AQP4 and LAMP1 signals (68 ± 12%; Fig. 2F), consistent with efficient targeting to lysosomal compartments. In contrast, wild-type AQP4 displayed reduced co-localization with LAMP1 (Fig. 2G), with significantly lower overlap values (48 ± 15%; Fig. 2H), indicating diminished lysosomal routing.

Together, these data demonstrate efficient delivery and expression of recombinant AQP4 constructs in mDCs. Importantly, incorporation of the LAMP3 sequence effectively redirects AQP4 to the endo-lysosomal pathway, a key step for enhanced antigen processing. Based on these findings, and consistent with previous work^32^, AQP4-LAMP3 (AQP4*) was selected for subsequent experiments, particularly in light of the limited availability of patient-derived biological material.

### Assessing APC Peptide Processing of Recombinant Antigens

The endo-lysosomal localization of AQP4* in HEK293 cells suggests that in antigen-presenting cells (APCs), its peptides may be processed and presented *via* MHC-II to CD4^+^ T cells. This raises a critical question: do these processed peptides generate immune epitopes recognized by T cells of NMOSD patients, potentially driving their disease? Answering this is crucial for evaluating their use in reestablishing tolerance in these patients.

To investigate this, we adapted an *in vitro* Ag-specific T cell proliferation assay^24^ (see **Fig. S3**). In this assay, monocyte-derived matured DCs (mDCs) were transfected with mRNA encoding specific antigens, then co-cultured with autologous CD4^+^ and CD8^+^ T cells (Schematics in Fig. 3A). To enhance Ag-specific T cell responses, a second stimulation was performed one week later using transfected mDCs. Activated T cells were identified by induction of CD154 expression on the surface of antigen stimulated CD4^+^ T cells^40^.In addition, increased activation of CD8^+^ T cells was analyzed by expression of CD69^+^ after the second stimulation.

**Figure 3.**
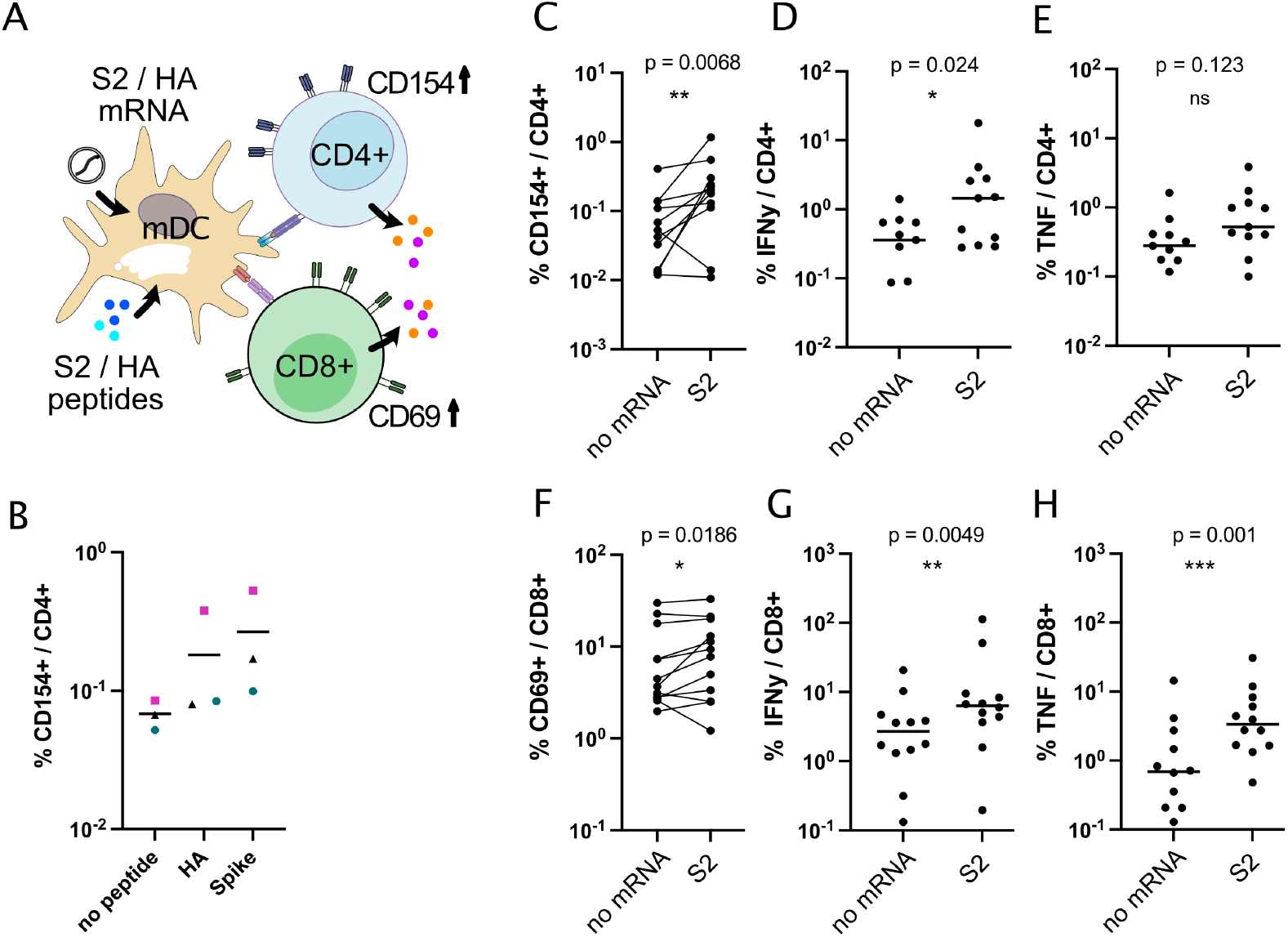
mRNA-loaded dendritic cells induce antigen-specific CD4^+^ and CD8^+^ T cell activation in healthy donors. (A) Schematic overview of antigen presentation following transfection of monocyte-derived dendritic cells (mDCs) with S2 or HA mRNA (Fig. S5). Translated proteins are processed into peptides and presented on MHC molecules, leading to activation of CD4^+^ T cells (CD154 upregulation) and CD8^+^ T cells (CD69 upregulation), accompanied by cytokine secretion. (B) Comparison of CD4^+^ T cell activation (%CD154^+^ within CD4^+^ cells) following stimulation with peptide pools or corresponding mRNA constructs, indicating comparable induction of antigen-specific responses. (C–E) CD4^+^ T-cell responses following stimulation with S2 mRNA. Activation (%CD154^+^/CD4^+^) (C) and IFNγ production (%IFNγ^+^/CD4^+^) (D) are significantly increased compared to control, whereas TNF production (%TNF^+^/CD4^+^) shows no significant difference (E). (F–H) CD8^+^ T-cell responses following S2 mRNA stimulation. Activation (%CD69^+^/CD8^+^) (F), IFNγ production (%IFNγ^+^/CD8^+^) (G), and TNF production (%TNF^+^/CD8^+^) (H) are significantly increased compared to control. Each dot represents an individual donor; lines indicate paired measurements where applicable. Statistical significance was assessed using paired comparisons; ns, not significant; *P < 0.05; **P < 0.01; ***P < 0.001.

To validate the assay, we assessed whether peptides or mRNA encoding SARS-CoV-2 Spike protein (S2) and influenza hemagglutinin (HA) could elicit T cell activation in healthy individuals. mDCs primed with S2 and HA peptides induced robust CD4^+^ T cell responses, as evidenced by increased frequencies of CD154^+^ CD4^+^ T cells compared to unstimulated controls (Fig. 3B). Similarly, mRNA encoding S2 or HA, with expression confirmed by immunofluorescence (Supplementary Fig. 1), elicited significant CD4^+^ T cell activation following transfection into mDCs and co-culture with autologous T cells (Fig. 3 C). These antigen-specific CD4^+^ T cells further exhibited increased production of IFNγ and TNF compared to non-transfected controls (Fig. 3D, E).

In addition, S2 mRNA-transfected mDCs induced activation of cytotoxic CD8^+^ T cells, as reflected by increased frequencies of CD8^+^CD69^+^ cells, along with enhanced IFNγ and TNF production (Fig. 3F–H). The effect of HA-mRNA transfected mDCs (Fig. S4) was less pronounced than S2, providing a rationale why we continued using S2-mRNA as a control mRNA for antigen-specific activation of T cells in following experiments.

Together, these results demonstrate that the mRNA-based platform enables efficient antigen delivery and processing in mDCs, leading to robust activation of both CD4^+^ and CD8^+^ T-cell responses *in vitro*.

### Assessing Antigenicity of Recombinant AQP4-M23-LAMP3 (AQP4*)

Our positive T cell responses to *ex vivo* mDCs transfected with mRNAs encoding foreign antigens (HA and S2) align with findings that mDCs can present antigenic peptides derived from mRNA-encoded proteins ^18,45^.

An open question remains whether such assays can detect immune responses to peptides processed from endogenous antigens like AQP4. Subcellular localization data suggests this may be possible for AQP4 tagged with the Lamp3 endo-lysosomal targeting sequence, which facilitates lysosomal processing (Fig. 2A, B).

To test this, we co-cultured T cells from healthy donors with autologous mDCs transfected with AQP4*-encoding mRNA, comparing these to untransfected controls and S2-transfected mDCs (Fig. 4A). While S2 elicited strong T cell activation (Fig. 4B), AQP4* did not provoke a similar response, likely due to insufficient antigen processing, suppression by regulatory T cells, and/or most likely a lack of antigen-specific effector T cells due to early thymic selection, as expected in a healthy condition. Similar challenges have been noted in tumor antigen studies ^37,46^ and may be addressed by co-expressing or adding immune activators.

**Figure 4.**
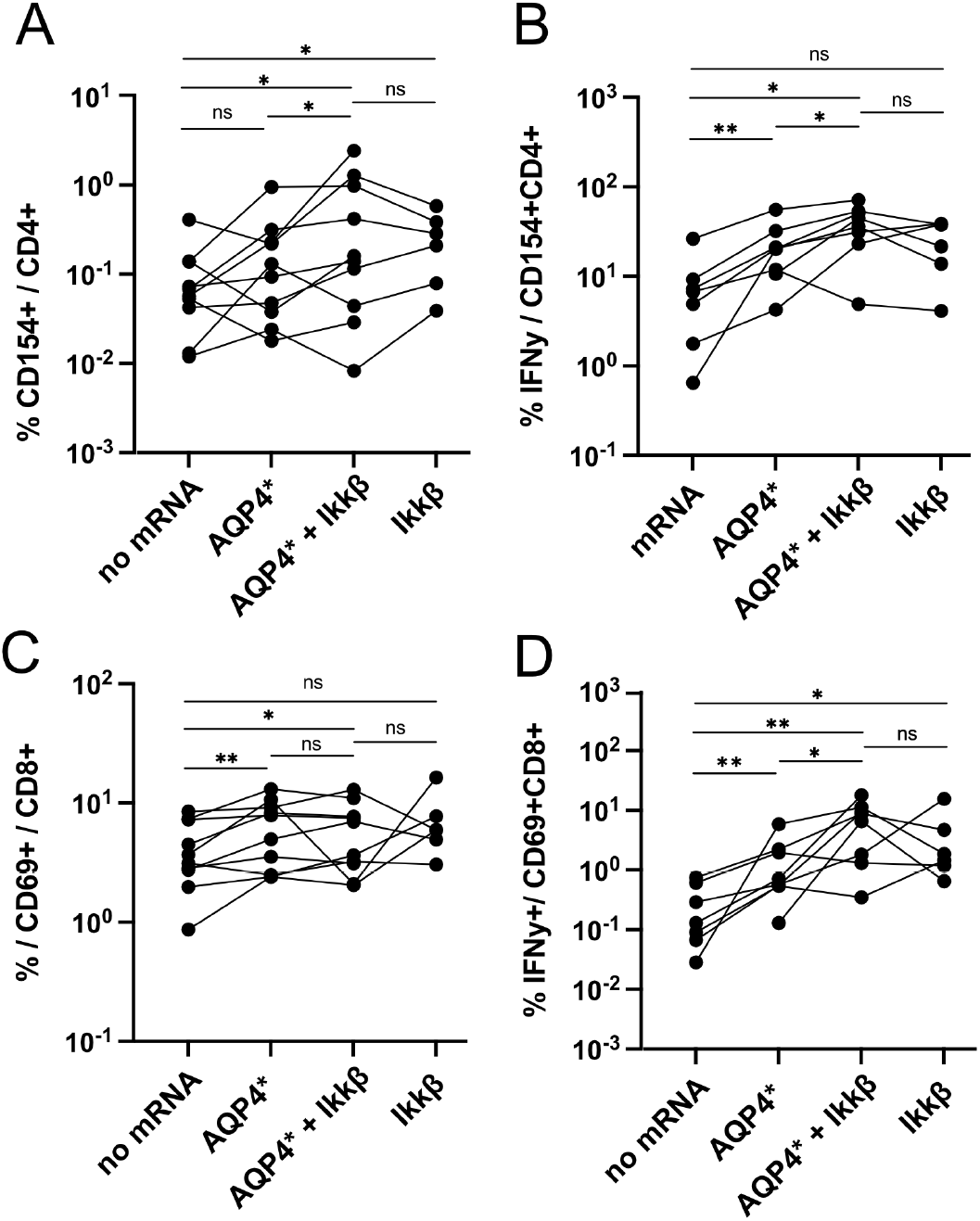
AQP4* mRNA delivery promotes CD4^+^ and CD8^+^ T cell activation and effector function. (A) Frequency of activated CD4^+^ T cells, measured as %CD154^+^ within CD4^+^ cells, following stimulation with no mRNA, AQP4*, AQP4* + IKKβ, or IKKβ alone. Paired donor responses show increased CD154 expression upon AQP4* delivery, with further modulation by co-expression of IKKβ. (B) IFNγ production in CD4^+^ T cells, expressed as %IFNγ^+^ within CD154^+^CD4^+^ cells. AQP4* induces cytokine production compared to control, with variable effects upon co-delivery of IKKβ. (C) Activation of CD8^+^ T cells assessed as %CD69^+^ within CD8^+^ cells. AQP4* stimulation increases CD8^+^ activation relative to no mRNA control, with no significant additive effect of IKKβ. (D) IFNγ production in CD8^+^ T cells, expressed as %IFNγ^+^ within CD69^+^CD8^+^ cells. AQP4* enhances effector responses, which are further increased in the presence of IKKβ in a subset of donors. Each line represents an individual donor. Statistical significance was assessed using Wilcoxon matched-paired comparisons; ns, not significant; *P < 0.05; **P < 0.01

Of note, there are a number of downstream effectors of NF-κB signaling, including TNF, IFNγ, CD83, CD80/86 that are involved in promoting immune responses by attracting T cells and enhancing the antigen-presentation ability of DCs ^38,47^. To enhance antigen-specific T cell activation, we generated a mRNA encoding constitutively active IKKβ, a key activator of NF-κB signaling, and transfected it either alone or in combination with AQP4* (Fig. 4A–D). IKKβ alone increased CD4^+^ T-cell activation, and co-transfection with AQP4* further augmented this response compared to AQP4* alone (Fig. 4A). This was accompanied by enhanced intracellular IFNγ accumulation in CD154^+^ CD4^+^ T cells, indicating increased functional activation upon co-delivery of IKKβ and AQP4* (Fig. 4B). In CD8^+^ T cells, AQP4* mRNA induced activation as reflected by upregulation of CD69, which was not further increased by co-expression of IKKβ (Fig. 4C). However, co-transfection with IKKβ resulted in elevated intracellular IFNγ levels, suggesting enhanced effector function within the CD8^+^ T cell compartment despite unchanged activation frequencies (Fig. 4D).

These results suggest that inducing an immune response against the self-antigen AQP4 is feasible, especially when immune activators like IKKβ are employed. This strategy may imitate mechanisms underlying AQP4 autoimmunity in patients ^48^.

### Exploring Ag-Specific T cell Responses in NMOSD Patients

We next investigated whether the Ag-specific T cell responses to recombinant AQP4* are enhanced in NMOSD patients. To evaluate the capacity of mRNA-loaded dendritic cells to induce antigen-specific T-cell responses, we stimulated autologous T cells with mDCs transfected with S2 or AQP4* mRNA (Fig. 5I).

**Figure 5.**
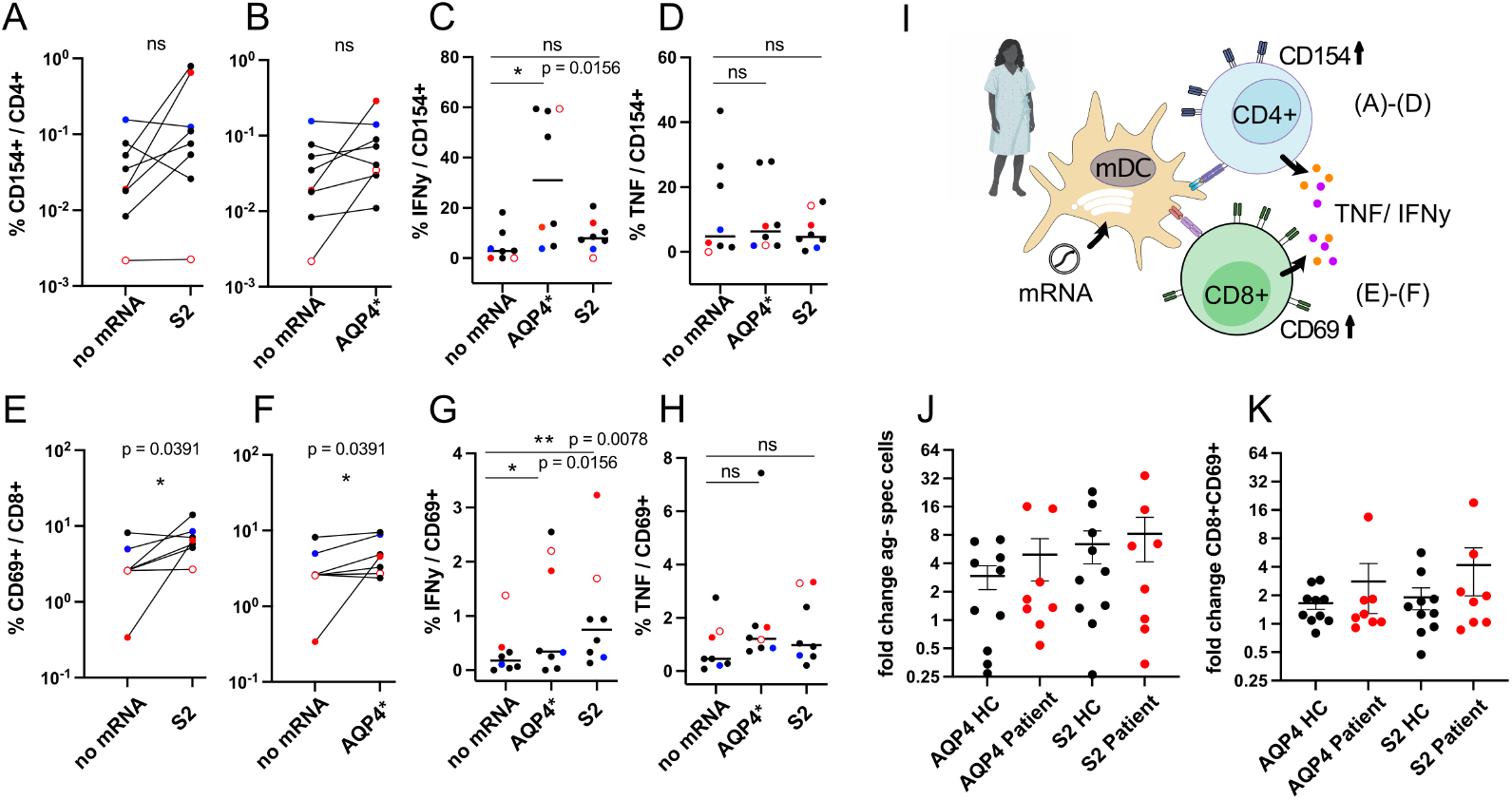
mRNA-driven antigen presentation induces CD4^+^ and CD8^+^ T cell responses in patients and healthy controls. (A, B) Activation of CD4^+^ T cells measured as %CD154^+^ within CD4^+^ cells following stimulation with S2 (A) or AQP4* (B) mRNA compared to no mRNA control. No significant differences were observed, red circle: patient 1, open red circle: patient 5, blue circle patient 2. (C, D) Functional profiling of CD4^+^ T cells. IFNγ production (%IFNγ^+^ within CD154^+^CD4^+^ cells) is increased following AQP4* stimulation compared to control (C), whereas TNF production (%TNF^+^ within CD154^+^CD4^+^ cells) remains unchanged across conditions (D). (E, F) Activation of CD8^+^ T cells assessed as %CD69^+^ within CD8^+^ cells following stimulation with S2 (E) or AQP4* (F) mRNA. Both conditions induce a modest but significant increase in CD8^+^ activation. (G, H) Effector function of CD8^+^ T cells. IFNγ production (%IFNγ^+^ within CD69^+^CD8^+^ cells) is significantly increased following stimulation, whereas TNF production (%TNF^+^ within CD69^+^CD8^+^ cells) is not significantly altered. (I) Schematic overview of the assay, illustrating mRNA-loaded monocyte-derived dendritic cells (mDCs) presenting antigen to CD4^+^ and CD8^+^ T cells, leading to activation (CD154, CD69) and cytokine production (IFNγ, TNF). (J, K) Comparison of antigen-specific responses between healthy controls (HC) and patients. Fold change in antigen-specific CD4^+^ T cells (J) and CD8^+^CD69^+^ T cells (K) is shown for AQP4 and S2 conditions. Individual data points are displayed with group means± SEM. Each dot represents an individual donor or patient. Statistical significance was assessed using Wilcoxon matched-paired comparisons; ns, not significant; *P < 0.05; **P < 0.01.

In NMOSD patients, S2 mRNA induced a trend toward increased CD4^+^ T cell activation, as measured by CD154 expression (Fig. 5A). Similarly, AQP4* mRNA did not significantly increase the frequency of CD154^+^ CD4^+^ T cells (Fig. 5B). However, functional analysis revealed a significant increase in IFNγ production within CD154^+^ CD4^+^ T cells upon AQP4* stimulation compared to unstimulated controls (Fig. 5C), while TNF production remained unchanged (Fig. 5D).

In the CD8^+^ compartment, both S2 and AQP4* mRNA induced significant activation, as reflected by increased frequencies of CD69^+^ CD8^+^ T cells (Fig. 5E, F). Moreover, IFNγ production in CD69^+^ CD8^+^ T cells was significantly enhanced following stimulation, with a more pronounced effect observed for S2 (Fig. 5G). As observed for CD154^+^ CD4^+^ T cells, TNF production in CD8^+^ T cells was also not significantly altered under these conditions (Fig. 5H).

To assess the translational relevance of these findings, we compared Ag-specific responses between healthy controls and patients. Fold-change analysis revealed comparable induction of antigen-specific CD4^+^ T cells in both groups following AQP4* and S2 stimulation (Fig. 5J) although a modest increased trend is visible for the patient group. Similarly, CD8^+^ T cell activation, assessed as fold change in CD69^+^ cells, showed no major differences between healthy donors and patients though a trend for the patient group (Fig. 5K).

Analysis of individual patient responses revealed marked heterogeneity in AQP4*-induced activation of antigen-specific CD4^+^ T cells. Notably, the strongest activation was observed in patient 1, who had experienced a recent relapse (2021), exhibited the highest AQP4 autoantibody titer (1:320), and was not receiving immunomodulatory therapy. In contrast, patient 2 showed no detectable response to either AQP4* or S2 stimulation; this patient had been relapse-free for over 10 years and lacked measurable AQP4 autoantibodies, despite also not receiving immunomodulatory treatment (Table 1). A pronounced response to AQP4* was also observed in patient 5, who presented with advanced disease progression/ high EDSS score, suggesting that disease stage may further influence T cell responsiveness.

Importantly, all patient samples responded robustly to the superantigen Staphylococcal Enterotoxin B (SEB), confirming preserved T cell functionality and validating the experimental system (Fig. S6).

Collectively, these findings suggest that immunomodulatory therapy, commonly administered in NMOSD patients, dampens AQP4-specific T-cell responses and may reduce or eliminate detectable antigen-specific T and B cell populations, even under *in vitro* stimulation conditions. Conversely, recent disease activity, high AQP4 autoantibody titers, and the absence of immunosuppressive treatment appear to favor robust antigen-specific T cell activation, as exemplified by patient 1. Patients with prolonged remission or under immunosuppressive therapy displayed reduced responsiveness.

Despite this variability, a significant increase in IFNγ production was observed in antigen-specific CD4^+^ (Fig. 5C) and CD8^+^ T cells (Fig. 5G) in several patients, indicating a persistent pro-inflammatory potential even under immunosuppressive conditions. The recognition of AQP4*-derived peptides by patient T cells further supports their relevance as potential biomarkers of disease activity in NMOSD.

Finally, the consistent SEB responsiveness across all samples underscores the functional competence of patient-derived immune cells and supports the robustness of the assay (Fig. S6).

## Discussion

In this study, we developed a system wherein antigen-presenting cells (APCs) were used to mimic the natural processing mechanism from mRNA to peptide and potentially display a broader, personalized repertoire of peptides bound by major histocompatibility complex (MHC-I or MHC-II). Engineered mRNAs expressing foreign antigens have revolutionized the development of vaccine therapies ^17,18,41^ that can be delivered subcutaneously, where antigen presenting Langerhans cells migrate to the lymph nodes boosting an immune response in a pro-inflammatory milieu. Peripherally circulating dendritic cells can also present foreign antigens, mounting a pro-inflammatory protective immune response regarding different pathogens. Importantly, these circulating DCs also hold the key to the maintenance of tolerance to the body’s own proteins, lipids carbohydrates and nucleic acids^16^. For individuals, the loss of tolerance to subsets of self-antigens has been causally linked to autoimmune disease ^49^. Current treatments include the use of broadly acting, non-disease-specific immuno-suppressants and immuno-modulators, which lead to numerous side effects and can cause serious adverse events, alongside unknown long-term complications and no potential for cure^13^.

In the current study, we have designed and evaluated recombinant versions of HA, S2 and AQP4 for the ability of their processed peptides to engage the immune system of both healthy individuals and AQP4-IgG+ NMOSD patients. Our experiments showed that these recombinant proteins are readily expressed in matured dendritic cells (mDCs). Importantly, appending AQP4 with the C-terminal tail of LAMP3 greatly increased its navigation into lysosomes (Fig. 2). In healthy donors, mRNAs expressing foreign antigens (HA and S2) elicited Ag-specific T cell responses, consistent with previous studies. Little or no response was detected following the transfection of mRNAs encoding AQP4-M23-LAMP3. Importantly, responsiveness to AQP4* was achieved when mDCs were co-transfected with a second mRNA encoding the immune activating protein IKKβ ^37^, suggesting processed peptides from AQP4* can induce an Ag-specific response. Furthermore, we successfully isolated monocytes from PBMCs of NMOSD patients and transformed them into mDCs. These mDCs were then transfected with our recombinant mRNAs encoding AQP4* and subsequently incubated with the patients’ autologous T cells. T cells from NMOSD patients currently on immunosuppressive or immune-modulating drugs or no therapy with long time remission (>10 years since last relapse) did not or did only sparsely respond to mDCs primed with mRNAs encoding either AQP4* or S2. However, a clear response could be detected in the patient with the most recent disease activity combined with absence of immunosuppressive/immunomodulatory therapy. In addition, this patient exhibited clear positive levels of AQP4 antibodies. Although the included number of patients per group is limited due to the rarity of the disease, these data support the concept that our recombinant AQP4* when processed by mDCs can generate peptides that are recognized by the immune systems of NMOSD patients and lead to an enhanced T cell response in patients with more recent relapses. We therefore propose that this assay could be utilized to identify disease-specific biomarkers (e.g., T cell responsiveness to the disease-specific antigen AQP4), possibly enabling the prediction of ongoing disease activity in the future.

Currently, despite promising data for NfL and GFAP there are no clinically validated biomarkers available to address the following issues: determining disease severity, monitoring disease activity, guiding treatment decisions, tracking treatment response, and predicting relapses ^50,51^. Our method could serve as a tool for monitoring AQP4-specific T cells in NMOSD patients and thus serve as a disease-specific biomarker that indicates disease activity defined as currently present activated effector T cell against the antigen AQP4 and might help to assess subclinical disease activity^52^. This would facilitate predictions about disease progression and inform the necessity for continued immunosuppressive therapy.

Two findings from our NMOSD patient cohort were unexpected. First, six of the eight individuals showed little or no detectable T-cell response following co-culture with primed autologous mDCs expressing mRNAs encoding either AQP4-LAMP3 or S2 (Fig. 5). Second, two patients (patient 1, red circle, and patient 5, open red circle; Fig. 5A, B) exhibited strong AQP4-specific responses. While patient 1 had not received immunosuppressive therapy, patient 5 had been treated with RTX but was already at an advanced disease stage. The persistence of a robust AQP4-directed response in patient 5 suggests that pro-inflammatory immune potential may be maintained despite immunosuppressive treatment.

A subsequent review of each patient’s history revealed that 4/6 patients having little or no T cell response against these antigens had received a B cell depletion therapy (Rituximab (RTX) or Inebilizumab) (see Table 1). B cells are APCs and influence the T cell response. Bar-Or et al. (2010) have shown the influence of B cell depletion mediated by RTX on T cells in patients suffering from Multiple Sclerosis. A diminished Th1 and Th17 response was observed *ex* and *in vivo* under B cell depletion ^53^. Furthermore, the CD154^+^CD4^+^ T cell population is decreased in Systemic Sclerosis (SSc) and Systemic Lupus Erythematosus (SLE) patients after RTX ^54,55^. Consequently, this could be one factor of the low immune response against our mRNA in NMOSD patients. Intriguingly, the patient (patient 1) with the strongest T cell response against both AQP4* and S2 in our *in vitro* assay had not received any immunomodulatory therapy since 2021, had the shortest time since her last relapse, and the highest AQP4 autoantibody titer. Of note, patient 2 who had a weak T cell response also had not received immunomodulatory therapy, and notably had her last relapse 10 years ago, which might explain why her T cell response was similar to that seen in healthy controls. Furthermore, this latter individual had no measurable AQP4 autoantibody titer at the time of our study. Patient 5 While more data are needed, these preliminary findings suggest two things: Firstly, that our recombinant AQP4* when processed by mDCs can generate peptides that are recognized by the immune systems of NMOSD patients. Secondly, that the immune system of NMOSD patients receiving immunomodulatory therapy is in a suppressed state and thus either unresponsive to our engineered AQP4* mRNA activation approach or exhausted, as suggested by Saggau et al, (2024)^56^.

### Limitations of the Study

There was significant variability in individual responses to recombinant antigens (HA, S2, and AQP4*), likely due to the complexity of immune responses, health status, vaccination history, and timing of blood collection ^57–59^. In healthy donors, responses to AQP4-M23 were low compared to foreign antigens like HA and S2, consistent with central tolerance mechanisms. While co-delivery of AQP4* mRNA with an IKK-β activator enhanced CD4^+^ T cell activation (Fig. 4), increasing the number of NMOSD donors would improve statistical significance. Additionally, longitudinal analysis of therapy-naïve patients during relapse and long-term follow-up during and after therapy would provide valuable insights, though patient scarcity (1–4 per 100,000) (11) poses challenges for sample acquisition and processing.

### Outlook and Conclusion

Our study investigates the use of recombinant AQP4*-mRNA, processed by myeloid-derived dendritic cells (mDCs), to generate immune peptides in NMOSD patients. Unlike previous strategies, which rely on whole PBMC populations, our approach uses purified mDCs for antigen presentation, offering several advantages: (a) it focuses on cells specifically responsible for antigen presentation, (b) it eliminates complex T cell interactions with other PBMC components, and (c) it ensures more controlled experimental conditions. Additionally, the intracellular processing of mRNA offers greater flexibility in generating a broader repertoire of antigenic peptides, compared to pre-processed peptides of fixed lengths. mRNA represents a revolutionary approach to T cell stimulation, offering advantages over traditional peptide-based strategies. It is cost-effective, rapidly producible, and capable of delivering multiple antigens within a single construct, transforming immunotherapy. This approach is crucial for reestablishing tolerance *via* tolerogenic dendritic cells (tolDCs), which must present a repertoire of immunogenic peptides that drive autoimmunity against AQP4. mRNA-based reprogramming of tolDCs shows promise for restoring antigen-specific tolerance across autoimmune diseases. Notably, a phase IIb clinical study demonstrated that DC-AQP4 peptide-loaded therapy was safe and tolerable, inducing an anti-inflammatory response (e.g., elevated IL-10) ^57^. However, it did not elicit a specific response to AQP4 or myelin-associated proteins, highlighting the limitations of peptide-based strategies, such as narrow peptide repertoires and rapid turnover ^57^.

These challenges can be addressed by using full-length recombinant molecules like AQP4*-mRNA, which can deliver multiple antigenic peptides and factors to *ex vivo* reprogrammed tolDCs, facilitating antigen-specific immune tolerance. This approach reduces unwanted side effects and adverse events. The feasibility of mRNA-based strategies is supported by a study by Krienke et al., showing that non-inflammatory mRNA encoding myelin oligodendrocyte glycoprotein (MOG) could suppress disease progression in mice, offering potential for treating multiple sclerosis ^58^.

Our study demonstrates the potential for of mRNA-transfected dendritic cells to monitor Ag-specific immune responses of NMOSD patients. Importantly, we observed a strong immune response in a patient with a shorter time to last relapse that was not receiving immunosuppressive therapy. These findings have significant implications for future research in the discovery of biomarkers to predict disease course and efficacy of immunosuppressive therapeutics in patients suffering from NMOSD. Ultimately, this approach might even be engineered to achieve restoration of immune tolerance in patients suffering from NMOSD as novel therapeutic approach.

## Acknowledgments and Sources Funding

We thank Stefan Frischbutter and Dietmar Schmitz for the intellectual and conceptual contributions, Anny Kretschmer and Maryam Mohamaddokht for the excellent technical support and we also thank the Flow Cytometry Core Facility at German Rheumatism Research Centre (DRFZ).

This study was financially supported by German Center for Neurodegenerative Diseases (DZNE) Experimental and Clinical Research Center (ECRC), Deutsche Forschungsgemeinschaft (DFG), projectnumber 441072869, SPARK BIH 2022 & 2023 – project „ REGAIN” and a grant of the Rosemarie und Thomas Minta – Gedenkstiftung and the Foundation DZNE Stiftung-Innovative Minds Program. Dr. Nadine Strempel is participant of the national Translational Tandem Program for Gene- and Cell-based Therapies (nTTP-GCT) – coordinated by the Biomedical Innovation Academy of the Berlin Institute of Health at Charité (BIH) and funded by the Federal Ministry of Research, Technology and Space (BMFTR). E.K. acknowledges the German Research Foundation (DFG) (KL 3278/2-1) grant, by the Helmholtz Association through program-oriented funding as well as by Humboldt Universität zu Berlin.

## Authorship Contribution Statement

NS, MH and CG conceptualized the project. NS and VP performed the experiments. NS, CG and VP were responsible for the data curation. EK was helping with the bioinformatics and data analysis. NS, CG, FP, MH and DS managed the project administration. VP wrote the original draft, while CG, NS, VP, MH, EK and SC contributed to writing, editing and revising the paper. MH recruited and consented patients.

## References

1. Jarius, S. et al. Contrasting disease patterns in seropositive and seronegative neuromyelitis optica: A multicentre study of 175 patients. J Neuroinflammation 9, 14 (2012).

2. Lennon, V. A. et al. A serum autoantibody marker of neuromyelitis optica: distinction from multiple sclerosis. Lancet 364, 2106–12 (2004).

3. Jarius S et al. Neuromyelitis optica. Nat Rev Dis Primers 6, 85 (2020).

4. Fujihara K et al. Neuromyelitis optica should be classified as an astrocytopathic disease rather than a demyelinating disease. Clinical and Experimental Neuroimmunology 3, 58–73 (2012).

5. Uzawa, A., Oertel, F. C., Mori, M., Paul, F. & Kuwabara, S. NMOSD and MOGAD: an evolving disease spectrum. Nat Rev Neurol 20, 602–619 (2024).

6. Pohl M et al. Pathogenic T cell responses against aquaporin 4. Acta Neuropathol 122, 21–34 (2011).

7. Pohl M et al. T cell-activation in neuromyelitis optica lesions plays a role in their formation. Acta Neuropathol Commun 1, 85 (2013).

8. Zeka B et al. Highly encephalitogenic aquaporin 4-specific T cells and NMO-IgG jointly orchestrate lesion location and tissue damage in the CNS. Acta Neuropathol 130, 783–98 (2015).

9. Kalluri SR et al. Functional characterization of aquaporin-4 specific T cells: towards a model for neuromyelitis optica. PLoS One 6, e16083 (2011).

10. Varrin-Doyer M et al. Aquaporin 4-specific T cells in neuromyelitis optica exhibit a Th17 bias and recognize Clostridium ABC transporter. Ann Neurol 72, 53–64 (2012).

11. Afzali AM et al. B cells orchestrate tolerance to the neuromyelitis optica autoantigen AQP4. Nature (2024).

12. Ren Z et al. Cross-immunoreactivity between bacterial aquaporin-Z and human aquaporin-4: potential relevance to neuromyelitis optica. J Immunol 189, 4602–11 (2012).

13. Kümpfel, T. et al. Update on the diagnosis and treatment of neuromyelitis optica spectrum disorders (NMOSD) – revised recommendations of the Neuromyelitis Optica Study Group (NEMOS). Part II: Attack therapy and long-term management. J Neurol 271, 141–176 (2024).

14. Derdelinckx J et al. Clinical and immunological control of experimental autoimmune encephalomyelitis by tolerogenic dendritic cells loaded with MOG-encoding mRNA. J Neuroinflammation 16, 167 (2019).

15. Banchereau J & Steinman RM. Dendritic cells and the control of immunity. Nature 392, 245–52 (1998).

16. Steinman RM, Hawiger D & Nussenzweig MC. Tolerogenic dendritic cells. Annu Rev Immunol 21, 685–711 (2003).

17. Baldin AV, Savvateeva LV, Bazhin AV, Zamyatnin AA & Jr. Dendritic Cells in Anticancer Vaccination: Rationale for Ex Vivo Loading or In Vivo Targeting. Cancers (Basel) 12, (2020).

18. Dörrie J, Schaft N, Schuler G & Schuler-Thurner B. Therapeutic Cancer Vaccination with Ex Vivo RNA-Transfected Dendritic Cells-An Update. Pharmaceutics 12, (2020).

19. Sahin U et al. Personalized RNA mutanome vaccines mobilize poly-specific therapeutic immunity against cancer. Nature 547, 222–6 (2017).

20. Mahnke K, Qian Y, Knop J & Enk AH. Induction of CD4+/CD25+ regulatory T cells by targeting of antigens to immature dendritic cells. Blood 101, 4862–9 (2003).

21. Idoyaga J et al. Specialized role of migratory dendritic cells in peripheral tolerance induction. J Clin Invest 123, 844–54 (2013).

22. Maksimow M, Miiluniemi M, Marttila-Ichihara F, Jalkanen S & Hänninen A. Antigen targeting to endosomal pathway in dendritic cell vaccination activates regulatory T cells and attenuates tumor immunity. Blood 108, 1298–305 (2006).

23. Korokhov N et al. High efficiency transduction of dendritic cells by adenoviral vectors targeted to DC-SIGN. Cancer Biol Ther 4, 289–94 (2005).

24. Cimen Bozkus C, Blazquez AB, Enokida T & Bhardwaj N. A T-cell-based immunogenicity protocol for evaluating human antigen-specific responses. STAR Protoc 2, 100758 (2021).

25. Sperber, P. S. et al. Berlin Registry of Neuroimmunological entities (BERLimmun): protocol of a prospective observational study. BMC Neurol 22, 479 (2022).

26. Wingerchuk DM et al. International consensus diagnostic criteria for neuromyelitis optica spectrum disorders. Neurology 85, 177–89 (2015).

27. Jarius S et al. Update on the diagnosis and treatment of neuromyelits optica spectrum disorders (NMOSD) - revised recommendations of the Neuromyelitis Optica Study Group (NEMOS). Part I: Diagnosis and differential diagnosis. J Neurol 270, 3341–68 (2023).

28. Jung, J. S. et al. Molecular characterization of an aquaporin cDNA from brain: candidate osmoreceptor and regulator of water balance. Proc Natl Acad Sci U S A 91, 13052–6 (1994).

29. Crane, J. M. et al. Binding Affinity and Specificity of Neuromyelitis Optica Autoantibodies to Aquaporin-4 M1/M23 Isoforms and Orthogonal Arrays. Journal of Biological Chemistry 286, 16516–16524 (2011).

30. Verkman AS, Rossi A & Crane JM. Live-cell imaging of aquaporin-4 supramolecular assembly and diffusion. Methods Enzymol 504, 341–54 (2012).

31. Crane, J. M., Van Hoek, A. N., Skach, W. R. & Verkman, A. S. Aquaporin-4 dynamics in orthogonal arrays in live cells visualized by quantum dot single particle tracking. Mol Biol Cell 19, 3369–78 (2008).

32. Bonehill A et al. Messenger RNA-electroporated dendritic cells presenting MAGE-A3 simultaneously in HLA class I and class II molecules. J Immunol 172, 6649–57 (2004).

33. de Saint-Vis B et al. A novel lysosome-associated membrane glycoprotein, DC-LAMP, induced upon DC maturation, is transiently expressed in MHC class II compartment. Immunity 9, 325–36 (1998).

34. Braun J et al. SARS-CoV-2-reactive T cells in healthy donors and patients with COVID-19. Nature 587, 270–4 (2020).

35. Chan JF et al. Genomic characterization of the 2019 novel human-pathogenic coronavirus isolated from a patient with atypical pneumonia after visiting Wuhan. Emerg Microbes Infect 9, 221–36 (2020).

36. Kibria KMK et al. A conserved subunit vaccine designed against SARS-CoV-2 variants showed evidence in neutralizing the virus. Appl Microbiol Biotechnol 106, 4091–114 (2022).

37. Pfeiffer IA et al. Triggering of NF-kappaB in cytokine-matured human DCs generates superior DCs for T-cell priming in cancer immunotherapy. Eur J Immunol 44, 3413–28 (2014).

38. Liu T, Zhang L, Joo D & Sun SC. NF-κB signaling in inflammation. Signal Transduct Target Ther 2, 17023–(2017).

39. Jonuleit H et al. Pro-inflammatory cytokines and prostaglandins induce maturation of potent immunostimulatory dendritic cells under fetal calf serum-free conditions. Eur J Immunol 27, 3135–42 (1997).

40. Bacher P & Scheffold A. Flow-cytometric analysis of rare antigen-specific T cells. Cytometry A 83, 692–701 (2013).

41. Kranz LM et al. Systemic RNA delivery to dendritic cells exploits antiviral defence for cancer immunotherapy. Nature 534, 396–401 (2016).

42. Furman CS et al. Aquaporin-4 square array assembly: opposing actions of M1 and M23 isoforms. Proc Natl Acad Sci U S A 100, 13609–14 (2003).

43. Jarzebska NT et al. Lipofection with Synthetic mRNA as a Simple Method for T-Cell Immunomonitoring. Viruses 13, (2021).

44. Strobel I et al. Human dendritic cells transfected with either RNA or DNA encoding influenza matrix protein M1 differ in their ability to stimulate cytotoxic T lymphocytes. Gene Ther 7, 2028–35 (2000).

45. Liang F et al. Efficient Targeting and Activation of Antigen-Presenting Cells In Vivo after Modified mRNA Vaccine Administration in Rhesus Macaques. Mol Ther 25, 2635–47 (2017).

46. Schuler G. Dendritic cells in cancer immunotherapy. Eur J Immunol 40, 2123–30 (2010).

47. Zinatizadeh MR et al. The Nuclear Factor Kappa B (NF-kB) signaling in cancer development and immune diseases. Genes Dis 8, 287–97 (2021).

48. Qiao H et al. Changes in the BTK/NF-κB signaling pathway and related cytokines in different stages of neuromyelitis optica spectrum disorders. Eur J Med Res 27, 96 (2022).

49. Bar-Or A et al. Restoring immune tolerance in neuromyelitis optica: Part II. Neurol Neuroimmunol Neuroinflamm 3, e277 (2016).

50. Schindler, P. et al. Glial fibrillary acidic protein as a biomarker in neuromyelitis optica spectrum disorder: a current review. Expert Review of Clinical Immunology 19, 71–91 (2023).

51. Aktas, O. et al. Serum neurofilament light chain levels at attack predict post-attack disability worsening and are mitigated by inebilizumab: analysis of four potential biomarkers in neuromyelitis optica spectrum disorder. J Neurol Neurosurg Psychiatry 94, 757–768 (2023).

52. Molazadeh, N., Filippatou, A. G., Vasileiou, E. S., Levy, M. & Sotirchos, E. S. Evidence for and against subclinical disease activity and progressive disease in MOG antibody disease and neuromyelitis optica spectrum disorder. Journal of Neuroimmunology 360, 577702 (2021).

53. Bar-Or A et al. Abnormal B-cell cytokine responses a trigger of T-cell-mediated disease in MS? Ann Neurol 67, 452–61 (2010).

54. Antonopoulos I et al. B cell depletion treatment decreases CD4+IL4+ and CD4+CD40L+ T cells in patients with systemic sclerosis. Rheumatol Int 39, 1889–98 (2019).

55. Sfikakis PP et al. Remission of proliferative lupus nephritis following B cell depletion therapy is preceded by down-regulation of the T cell costimulatory molecule CD40 ligand: an open-label trial. Arthritis Rheum 52, 501–13 (2005).

56. Saggau C et al. Autoantigen-specific CD4(+) T cells acquire an exhausted phenotype and persist in human antigen-specific autoimmune diseases. Immunity (2024).

57. Zubizarreta I et al. Immune tolerance in multiple sclerosis and neuromyelitis optica with peptide-loaded tolerogenic dendritic cells in a phase 1b trial. Proc Natl Acad Sci U S A 116, 8463–70 (2019).

58. Krienke C et al. A noninflammatory mRNA vaccine for treatment of experimental autoimmune encephalomyelitis. Science 371, 145–53 (2021).

59. Bacher P et al. Antigen-reactive T cell enrichment for direct, high-resolution analysis of the human naive and memory Th cell repertoire. J Immunol 190, 3967–76 (2013).

